# Larger hybrid clutch size could drive regional displacement of native *Iguana* populations across the Lesser Antilles

**DOI:** 10.1101/2025.07.31.667849

**Authors:** Matthijs P. van den Burg, Sarah A. van der Horn, P-A Åhlen, Lara Jansen, Kai Wulf, Misha Paauw, Adolphe O. Debrot

## Abstract

Native *Iguana* populations across the Caribbean Lesser Antilles are being replaced by non-native green iguanas (NNGI) once these establish themselves, leading to eventual extinction. Despite this knowledge, and much effort towards NNGI removal and prevention of incursions, the underlying mechanisms of the observed genetic swamping are poorly understood. Previous studies show that larger clutch size is a key reproductive trait of mainland as opposed to insular lizard populations. Higher clutch size in NNGI and their progeny could be a driver in the genetic swamping of insular *Iguana* populations. We collected data on snout-vent length, tail length, clutch size and genetics opportunistically during an eradication program of hybrid and NNGIs on the island of Saba, home to a native *Iguana Iguana* population. Data of native and hybrid iguanas from Saba were combined with similar published data from native insular and mainland *Iguana* populations. Generalized linear modeling revealed that climatic variables could not explain differences in clutch size, in contrast to genetic origins. Our preliminary results show that hybrid iguanas lay significantly more eggs (~40%) than similar-sized native Saba iguanas. Therefore, we suggest that higher hybrid clutch size may be a contributing factor hastening the genetic swamping and declines in native *Iguana* populations across the Lesser Antilles. Further study is needed to clarify and better understand the drivers of the rapid genetic swamping and loss of Antillean iguana diversity, which should also focus on *Iguana delicatissima* and utilize genomic approaches to pinpoint genes under selection.

## Introduction

Global biodiversity is threatened significantly by invasive alien species (IAS) that negatively impact native species through a range of processes like resource competition, introduction of diseases, predation, and hybridization (e.g., Allendorf et al. 2001; Strubbe and Matthysen 2009). Hybridization and introgression with IAS (i.e. anthropogenic hybridization; Allendorf et al. 2001) can have different diminishing impacts on native species (e.g., loss of genetic diversity, wasted reproductive effort, see Allendorf et al. 2001), and has even led to local extirpations (Rhymer & Simberloff 1996; Todesco et al. 2016). Conservationists are especially challenged when hybridization occurs between a rare species and closely-related IAS. Not only because this process can easily go undetected (e.g., Vanhaecke et al. 2012; Hata et al. 2019), but also since protecting the rare endangered species especially requires efforts to prevent hybridization. However, the mechanisms of decline due to hybridization remain poorly understood.

Native iguanas within the genus *Iguana* were once present across islands in the Caribbean Lesser Antilles, but have been either extirpated from numerous islands or have dramatically decreased in population size due to a range of direct and indirect anthropogenic causes (Knapp et al. 2014, 2021; Vuillaume et al. 2015; van den Burg et al. 2018a; Breuil et al. 2019). Within this insular region the current most severe threat to remaining native *Iguana* populations is the region-wide presence of non-native green iguanas (NNGI) (Breuil et al. 2019; Knapp et al. 2021; van den Burg et al. 2023). Their initial arrival to the region on the islands of Les Saintes dates to the 1860’s (Vuillaume et al. 2015), while more recently, natural (hurricanes, Censky et al. 1998) and unnatural (e.g., as stowaways in cargo containers) intra-regional translocations have occurred resulting in the establishment of non-native populations (van den Burg et al. 2018b, 2020).

Impacted native iguanas include *Iguana delicatissima* Laurenti, 1768 (from Anguilla to Martinique, except Saba and Montserrat) and members of the *Iguana Iguana* species complex (Linnaeus, 1758) (sensu Iguana Taxonomy Working Group, 2016). Considering the entire *I. iguana* complex, these populations belong to one of the four major mtDNA clades (named clade IIb in Stephen et al. 2013; *I. iguana* in van den Burg et al. 2021; clade IV van den Burg et al. 2026). Following a number of recent manuscripts on the taxonomy of these insular Lesser Antillean *I. iguana* populations (Breuil et al. 2019, 2020, 2022), currently the subspecies of *I. iguana insularis* (St. Vincent and the Grenadines and Grenada) and *I. iguana sanctaluciae* (St. Lucia) are recognized, but not the species *I. melanoderma* (also not recognized as subspecies; Saba and Montserrat) (Iguana Taxonomy Working Group, 2022; van den Burg et al. 2026). Additionally, the Iguana Taxonomy Working Group (2022) currently recognizes *I. iguana iguana* and *I. iguana rhinolopha*; see geographic limitation for *I. i. rhinolopha* in van den Burg et al. (2021a).

Non-native green iguanas within the Lesser Antilles have a diverse origin, which have been traced through genetic assessments of NNGIs from several islands using geographic referenced datasets from the native range (see van den Burg et al. 2026 for clade codes). These studies indicate origins from all four mtDNA clades, especially from Central America (Clade III) and northwestern South America (Clade II), with only few non-native populations including iguanas with (partial) origins from eastern South America (Clade IV) and the Curaçao clade (Clade I) (Vuillaume et al. 2015; van den Burg et al. 2018b, 2023, 2024; Pounder et al. 2020). However, a complete overview is still lacking, as several islands lack assessments or include only limited analyzed samples.

The arrival of NNGIs to Lesser Antillean islands has repeatedly led to anthropogenic hybridization and introgression with native *Iguana* populations (Day and Thorpe 1996; Vuillaume et al. 2015; van den Burg et al. 2018a; Breuil et al. 2019). Since this process leads to genetic swamping, including complete admixture without selection against hybrids with eventual extinction of the native population (Allendorf et al. 2001), much effort is being invested, through morphological (Breuil 2013; van den Burg et al. 2023, 2024) and genetic methods (Day and Thorpe 1996; Valette et al. 2013; Pounder et al. 2020; van den Burg et al. 2021a; van Kuijk et al. 2025), to rapidly identify and remove NNGIs. However, the mechanisms behind genetic swamping have received less research attention and are mostly limited to anecdotal data from *I. delicatissima* populations invaded by NNGIs. Vuillaume et al. (2015) indicate that hybrid and introgressed iguanas are “bigger and longer” than *I. delicatissima* and non-native males are more “powerful,” but without providing data. Using morphometric data, van den Burg et al. (2024) showed that NNGI on St. Maarten/Martin attain larger overall size compared to *I. delicatissima* on St. Eustatius, which also holds for native green iguanas from Saba (*I. i. iguana*; van den Burg et al. 2026), but a broader geographic study is needed to confirm this as there is variation in both parental species (Day and Thorpe 1996; Bock et al. 2018). Whether native X non-native hybrids also attain larger overall size compared to native iguanas has remained unassessed. Additionally, Vuillaume et al. (2015) suggested that the difference in reproductive timing between native and non-native iguanas could also play a role.

Clutch size has long been recognized as a key life-history trait affecting fitness and competitive ability (Godfray et al. 1991). While clutch size is often also correlated to other variables like egg weight and hatchling mass, these relationships are largely overrated, even in bird guilds where the relationship between clutch and egg parameters has been most intensively studied (Rohwer 1988). Thus while clutch size has several other correlates of significance like egg size and brood frequency, the key difference in reproductive traits between mainland and insular lizard populations is the difference in size-specific clutch size (Novosolov and Meiri 2013; Meiri et al. 2020). Hence, considering the observed genetic swamping of native island *Iguana* populations and thus a decline in future population size, clutch size is an important life-history trait to understand, as it directly reflects reproductive output and can influence future population growth. Especially, insights on potential differences between native and non-native iguanas and their hybrids are of great interest to understand future population trends and potential implications of biological invasions. Whilst it is known that *I. delicatissima* lays fewer eggs than (most of the) populations in the *I. iguana* species complex (Knapp et al. 2016; Bock et al. 2018). Very little data is available from native populations of the *I. iguana* species complex from islands in the Lesser Antilles (but see Blankenship 1990). Regarding hybridization, only a single published record of clutch size for an F1 *I. delicatissima* x *I. iguana* that suggests a larger clutch size in hybrids compared to *I. delicatissima* (see van Wagensveld and van den Burg 2018 and references therein). To contribute to a better understand the underlying mechanism(s) of the regional competitive displacement of native insular *Iguana* populations, we studied the difference in clutch size between native, invasive and hybrid *I. i. iguana* on the Lesser Antillean island of Saba.

### Materials and Methods

On Saba, the presence of NNGIs was first identified in 2021 (van den Burg et al. 2025a), which catalyzed a program to map their spread and remove as many non-native individuals as possible. Efforts within this program were sporadic during 2022 and 2023, but became more structural with a major NNGI removal effort during the reproductive season between December 2024 and March 2025. During these efforts we surveyed for and culled pure non-native and hybrid iguanas following morphological assignment based on the presence (native assignment)/absence (non-native or hybrid assignment) of the melanistic lateral spot between the eye and the tympanum (van den Burg et al. 2025a), the relative length of the head and dorsal spines and presence of enlarged nasal scales (van den Burg et al. 2023; van den Burg et al. 2025b). Continuous genetic assessment of removed iguanas indicates that only a minor percentage >5% was incorrectly identified (van den Burg et al. 2023, 2025b, unpublished data), and that these can correctly be identified ater they become roughly two years old. We also opportunistically captured native iguanas to collect morphological data, which were thereater released at their capture location. Additionally, we culled a few native females given they were part of a harem that included a large non-native dominant male and the high probability that mating had already taken place. Prior to December 2024, iguanas were either culled using an air rifle or through euthanasia ater noose/hand captures (lethal injection of pentobarbital by veterinarian), whilst most iguanas were shot using either a .22 or using different types of high-caliber ammunition during the 2024-2025 culling effort; .223, .243 and 6.5X55 with sot point ammunition for longer distances). This change of methodology was due to e?ciency since approaching iguanas for close-proximity removal can often result in them hiding. At longer distances, and given windy conditions on Saba, accurate bullet placement was aided using the Vortex Fury HD 5000 AB distance measuring feature and the Fury HD Vortex application, and all shots preluded a thorough assessment of backstop presence and orientation.

This culling effort allowed us to collect data on snout-vent length (SVL; in mm), tail length (TL; in mm), and clutch size, except for those bodies that could not be retrieved or when these variables were affected by bullet impact. From each captured or retrieved iguana we also sampled blood (using non-heparinized syringe) or tissue (using single-use scalpel), which were stored in Longmire buffer (Longmire et al. 1992) or >95% ethanol, respectively. Thereafter, we performed necropsies on all females to determine reproductive state and count the number of developing eggs (>1 cm diameter) to score clutch size. When SVL or the number of eggs was affected by bullet impact we excluded the iguana from our dataset (*n* = 3). Lastly, prior to 2024, we collected SVL and clutch size data opportunistically from a single native female that was hit by a car in 2021 (van den Burg et al. 2025). Fieldwork permits were obtained through the Public Entity Saba (permits BC663.21, BC515.2023, and BC428.2024), and we worked following recommendations from the IUCN SSC Iguana Specialist Group (IUCN SSC Iguana Specialist Group 2017) and euthanasia guidelines from the American Veterinary Medical Association.

We constructed two datasets to assess differences in total size (SVL + TL from animals with complete and unbroken tails) and clutch size among different groups. We assessed differences in total size among native iguanas from Saba (*n* = 25), NNGIs from St. Maarten (*n* = 22, van den Burg et al. 2024, 2025b), and genetically-confirmed hybrid iguanas from Saba (*n* = 17; for genetics see below and Appendix 1). Hereafter we retrieved residuals from a SVL-tail length regression, and compared the means of these SVL-corrected tail length differences using an analysis of variance (ANOVA) including subsequent posthoc Tukey test to assess different results between the groups. Considering clutch size, we assessed SVL-clutch size relationships among native and non-native iguanas from Saba, as well as previously published data from an *I. i. iguana* population in Panama (Rand 1984; Miller 1987), an *I. i. rhinolopha* population Nicaragua (Fitch & Henderson 1977), one population of *I. delicatissima* (Knapp et al. 2016), and the only reported *I. delicatissima* x *I. iguana* F1 hybrid datapoint (van Wagensveld & van den Burg 2018). First, we assessed data normality and homogeneity of variances using a Shapiro-Wilk and Levene’s test, respectively. We then assessed differences in regression slopes for the SVL – clutch size relation among these groups (excluding the F1 hybrid record) through a linear model and pairwise comparisons of regression coe?cients and least-square means. Then, for groups with equal slopes, we used generalized linear modeling (GLM) to assess whether differences in SVL and clutch size among these groups are explained by genetic/population status or climatic variables. We ran GLM analyses on two datasets, both with clutch size as the response variable. For the first analysis, we included the predictor variables SVL and genetic status, whilst for the second analysis we also included the WorldClim variables. We used three bioclimatic variables from the WorldClim 2 dataset (Fick & Hijmans 2017) of importance considering variation in reptilian clutch sizes (BIO1, annual mean temperature; BIO4, temperature seasonality; BIO15, precipitation seasonality; Meiri et al. 2020). Since clutch size is count data we used Poisson models. For this, we used analysis of deviance (AoD) (with Chisquare-based estimate) to assess relations between the response and predictor variables, and individual coe?cient results for the relations of between the response and categories within the predictor variables. Both models (with and without the bioclimatic variables) were compared using the Akaike information criterion (AIC). We used visual assessments of residuals, quantile-quantile plot and Cook’s statistics to assess model normality, assumptions and disproportionality. In case bioclimatic variables had no effect on clutch size difference, we tested for differences in SVL-corrected clutch size among the different genetic groups using an overall ANOVA and individual *t*-tests between all combinations; *p*-values were corrected using the BY-FDR method (White et al. 2019).

Blood/tissue samples were used to assess species/hybrid status for each female iguana and all iguanas with complete SVL+tail data, following van den Burg et al. (2023, 2024). Ater DNA extraction using a DNeasy Kit (QIAGEN, Germany), we performed Sanger sequencing (forward and reverse) at the University of Amsterdam where laboratory methods were implemented following Malone et al. (2017) and van den Burg et al. (2018b) to amplify the mitochondrial NADH dehydrogenase subunit 4 (ND4) and the nuclear MutL homolog 3 (MLH3). Using Geneious Prime 2024.0.4 (Kearse et al. 2012), we quality-checked chromatograms and compared consensus sequences per individual to published GenBank records and unpublished sequences. Additionally, for the female iguanas only, we generated microsatellite data for 17 loci commonly used for *Iguana* (Valette et al. 2013; Vuillaume et al. 2015; van den Burg et al. 2021a) at the Qualyse laboratory in France. Generated allelic data were compared to the IguanaBase database (van den Burg et al. 2021b) using discriminant analyses of principal components (DAPC) using the *adegenet* package (Jombart 2008) in the R environment (R Core Team 2025, version 4.3.2; RStudio Team 2025, version 2023.12.1+402). Since very few genetic data are available from St. Maarten/Martin, we were not able to employ a program to analytically predict the status of hybrid individuals (F1/F2), which we instead tentatively did using the mtDNA and nDNA haplotypes.

## Results

For clutch size comparisons, we obtained useful data from culled gravid female iguanas during Dec 2024–March 2025, indicating that both native, hybrid and non-native iguanas are reproductively active in the same period (Table 1). The variables SVL and clutch size had normal distributions (SVL, *p* = 0.059; eggs, *p* = 0.691) and equal variances (SVL, *p* = 0.245; eggs, *p* = 0.114).

**Table 1.** Collected data on snout-vent length (SVL) and egg count from female iguanas on Saba, including status assignment that was based on the combined morphological and genetic assignment for mitochondrial (mtDNA), nuclear (nDNA), and microsatellite (msat) data. Unless native, clade numbers in the assignment columns refer to mtDNA clades within the *Iguana Iguana* species complex following van den Burg et al. (2026). Asterisk: SAB88 was not included in further analyses since it lacked native haplotypes. Superscripted B: tails that were not complete.

| Individual | SVL (mm) | Tail (mm) | Eggs | Assignment | Morphology | mtDNA | nDNA | msat |
| --- | --- | --- | --- | --- | --- | --- | --- | --- |
| SAB45 | 325 | NA | 29 | native | native | native | native / native | native |
| SAB103 | 305 | 450<sup>B</sup> | 21 | native | native | native | native / native | NA |
| SAB104 | 345 | 500<sup>B</sup> | 29 | native | native | native | native / native | NA |
| SAB108 | 354 | 701<sup>B</sup> | 34 | native | native | native | native / native | native |
| SAB114 | 381 | 955 | 44 | native | native | native | native / native | native |
| SAB97 | 357 | 811 | 42 | F1 hybrid | hybrid | Clade III: HM352508 | native / Clade III: OQ556118 | nonnative |
| SAB100 | 369 | 921 | 50 | F1 hybrid | hybrid | native | native / Clade I: KX610637 | nonnative |
| SAB102 | 340 | 880 | 45 | F1 hybrid | hybrid | native | native / Clade III: PV974928 | nonnative |
| SAB117 | 345 | 843 | 42 | F1 hybrid | hybrid | native | native / Clade III: OQ556117 | nonnative |
| SAB80 | 361 | 875 | 40 | F1 hybrid | nonnative | native | native / Clade III: PV974928 | nonnative |
| SAB101 | 379 | 855<sup>B</sup> | 58 | F1 hybrid | nonnative | Clade III: HM352508 | native / Clade III: OQ556117 | nonnative |
| SAB91 | 311 | 802 | 37 | F2 hybrid | hybrid | Clade III: HM352508 | native / native | nonnative |
| SAB88* | 371 | 901 | 48 | nonnative | nonnative | Clade II: PV974931 | Clade III: OQ556117 / Clade III: PV974930 | nonnative |

We obtained ND4 and MLH3 sequences data from all 13 gravid females (Table 1); five individuals only had haplotypes native to Saba (ND4 and MLH3 haplotypes correspond to HM352505 and Oti556119, respectively), seven tentative hybrid individuals had native haplotype(s) and one or two haplotypes of Clade I or III, and one individual only had haplotypes of Clade II (mtDNA, PV974931) and III (nDNA, Oti556117 and PV974930). Out of the seven tentative hybrids, all except SAB91 show mtDNA/nDNA signs of F1 hybrids, with SAB91 being either a F2 hybrid or backcross. Considering non-native haplotype origin, the two unique mtDNA haplotypes originated from Clade II (PV974931) and III (HM352508), and the four unique nDNA haplotypes from Clade I (KX610637) and III (OQ556117, OQ556118, PV974928, and PV974930) (Table 1).

We also generated microsatellite allelic data from all but two samples (SAB103+104; Figure 1). DAPC clustering for these microsatellite alleles showed that samples either clustered within “*Iguana iguana* melanistic” (which were considered native given previous assessments; van den Burg et al. 2021a, 2023), or were placed outside of the “*Iguana iguana* melanistic” cluster. These later samples showed alleles not observed in native Saba iguanas (Breuil et al. 2020; van den Burg et al. 2021a, 2023), and were thus considered non-native (Table 1). Combined, for the 13 gravid female iguanas, these genetic data indicated that five were native Saba iguanas (SAB45, 103, 104, 108, 114), six were native X non-native F1 hybrids (SAB80, 97, 100, 101, 102, 117), and one iguana was a F2 hybrid (SAB91). SAB88 was a non-native iguana with mixed geographic origins and was therefore not included in SVL-clutch size comparisons.

**Figure 1.**
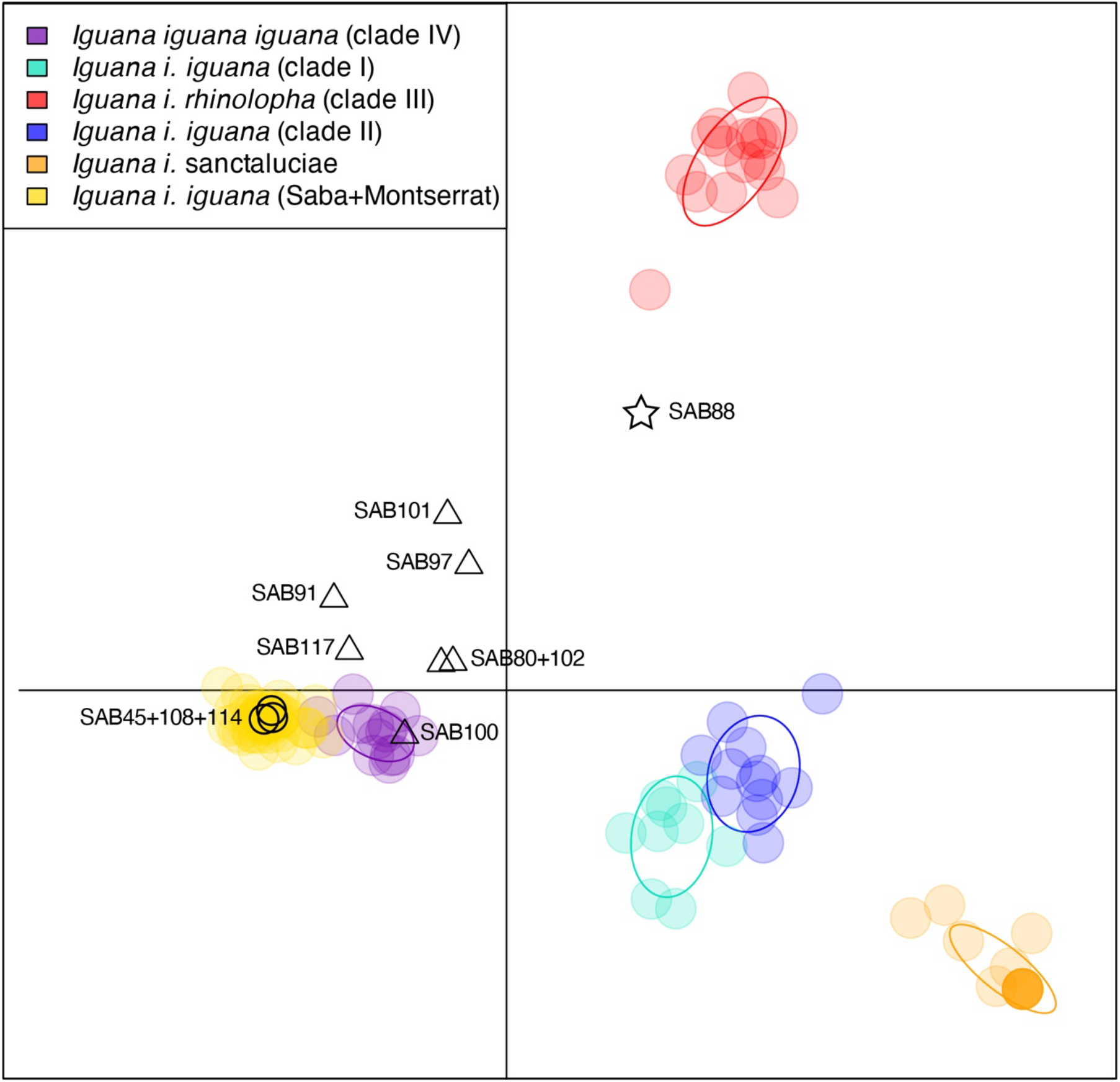
Discriminant analysis of a principal components scatterplot representing 17 microsatellite loci from reference IguanaBase individuals (colored circles) and 11 gravid females from Saba; three native iguanas (circles), and eight hybrid (triangles) and nonnative (star) iguanas. Clade codes follow van den Burg et al. (2026).

SVL-clutch size regression slopes differed significantly only between *I. delica,ssima* – *I. i. iguana* Panama (*p* = 0.0003) and *I. delica,ssima* – *I. i. rhinolopha* Nicaragua (*p* = 0.0302) (Figure 2A). Therefore, *I. delica,ssima* (as well as the single *I. delica,ssima* x *I. i. iguana* record) was not included in the generalized linear modeling datasets. Results of both GLM analyses indicated a significant effect of SVL on clutch size of native Saba iguanas (Table 2 and 3). For the model excluding bioclimatic variables (AIC score of 517.66), the analysis of deviance indicated a significant effect of SVL (*p* < 2.2^e-16^) and Status (*p* = 0.00047) on clutch size. Specifically, we found a significant effect of SVL on clutch size of native Saba iguanas, and a significant relation between clutch size of native Saba iguanas and hybrid iguanas on Saba (Table 2). Post-hoc analyses of pairwise comparisons between clutch sizes of the different groups (Saba native, Saba hybrid, Panama, and Nicaragua) showed that clutch size of hybrid iguanas on Saba was significantly larger than native Saba iguanas (with Tukey correction = 0.752, SE = 0.074, *p* = 0.018) and iguanas from Nicaragua (with Tukey correction = 1.282, SE = 0.084, *p* = 0.001), and clutch size of iguanas from Panama was significantly larger than iguanas from Nicaragua (with Tukey correction = 0.893, SE = 0.038, *p* = 0.043). Thus an average native female of 33.75 SVL females had an average clutch size of 30 eggs whereas F1 hybrids had an average clutch size of 42 eggs (40% higher) (Figure 2A).

**Table 2.**
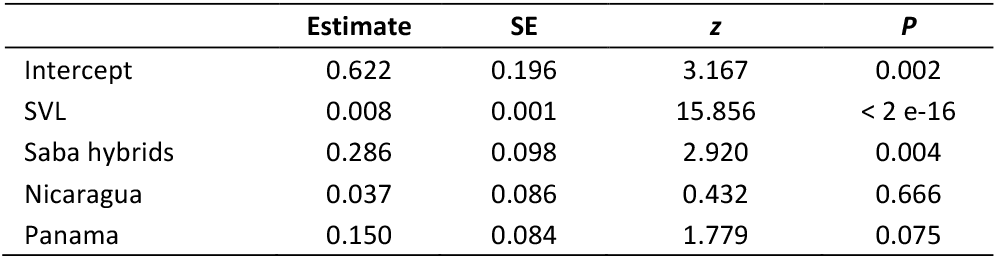
Results of the generalized linear mixed model (GLM) for factors affecting clutch size of native Saba iguanas, dataset excluding bioclimatic variables.

|  | Estimate | SE | z | P |
| --- | --- | --- | --- | --- |
| Intercept | 0.622 | 0.196 | 3.167 | 0.002 |
| SVL | 0.008 | 0.001 | 15.856 | < 2 e-16 |
| Saba hybrids | 0.286 | 0.098 | 2.920 | 0.004 |
| Nicaragua | 0.037 | 0.086 | 0.432 | 0.666 |
| Panama | 0.150 | 0.084 | 1.779 | 0.075 |

**Table 3.** Results of the generalized linear mixed model (GLM) for factors affecting clutch size of native Saba iguanas, dataset including bioclimatic variables; BIO1, annual mean temperature; BIO4, temperature seasonality; BIO15, precipitation seasonality (Fick & Hijmans 2017).

|  | Estimate | SE | z | P |
| --- | --- | --- | --- | --- |
| Intercept | -0.762 | 8.042 | -0.095 | 0.925 |
| SVL | 0.008 | 0.001 | 15.698 | < 2 e-16 |
| Saba hybrids | 0.316 | 0.108 | 2.914 | 0.004 |
| Nicaragua | -3.365 | 21.510 | -0.156 | 0.876 |
| Panama | -1.305 | 11.982 | -0.109 | 0.913 |
| BIO1 | -0.057 | 0.648 | -0.087 | 0.930 |
| BIO4 | 0.008 | 0.030 | 0.264 | 0.792 |
| BIO15 | 0.055 | 0.333 | 0.165 | 0.869 |

**Figure 2.**
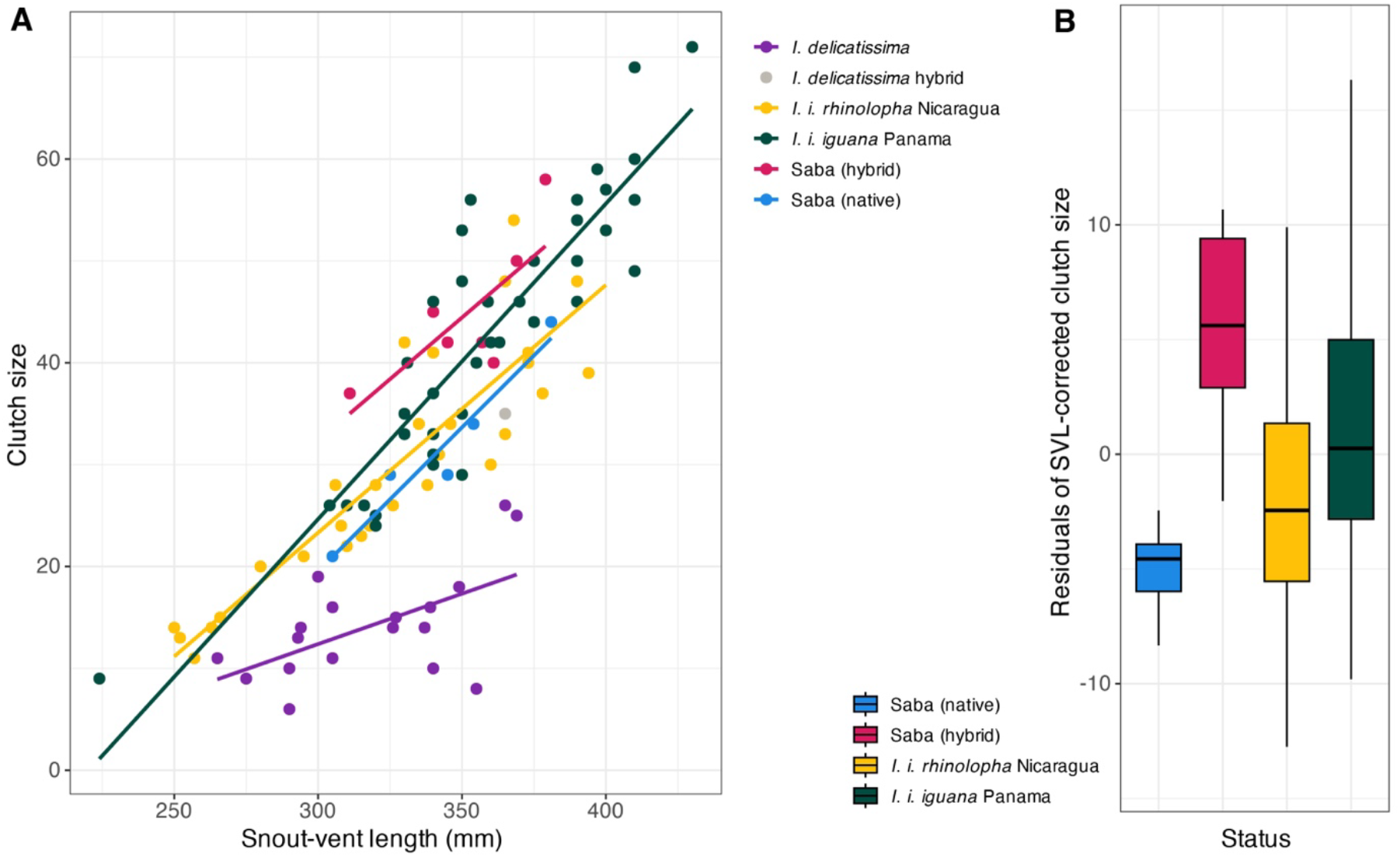
Relationships between snout-vent length (SVL) and clutch size among native and hybrid iguanas on Saba and different populations of *Iguana* spp., including data from Fitch & Henderson (1977), Rand (1984), Miller (1987), Knapp et al. (2016), and van Wagensveld & van den Burg (2018). **A)** SVL-clutch size regression including all groups. **B)** Boxplots of SVL-clutch size residuals among groups with regression slopes that were not significantly different. SAB88 is not included.

For the model including bioclimatic variables, the analysis of deviance indicated a significant effect of SVL (*p* < 2.2^e-16^) and Status (*p* = 0.00047) on clutch size, and no effect of BIO1 (*p* = 0.607), BIO4 (*p* = 0.526), and BIO15 (*p* = 0.869). Results of the GLM coe?cients were also not significant for the bioclimatic variables (Table 2). The model including the bioclimatic variables had an AIC score of 522.99, higher than the model excluding the bioclimatic variables. Visual assessments of residuals, quantile-quantile plot and Cook’s statistics showed no clear deviations of model normality, assumptions and disproportionality of observations (Supplementary figures 1 and 2). Since the model excluding the bioclimatic variables had a lower AIC, we used an ANOVA to assess the pairwise comparisons of SVL-corrected clutch size. The ANOVA indicated significant differences among SVL-corrected clutch size among means of the four groups (Figure 2B), *p* = 0.0011; Saba (native), Saba (hybrid), *I. i. rhinolopha* Nicaragua, and *I. i. iguana* Panama. HY-FDR adjusted *p*-values for pairwise comparisons were significant for all but two comparisons (Table 4): Saba (native) – *I. i. rhinolopha* Nicaragua and Saba (hybrid) – *I. i. iguana* Panama.

**Table 4.** Results of pairwise *t*-test for differences in SVL-corrected residuals in clutch size for female iguanas across four groups; Saba (native), Saba (nonnative), *I. i. rhinolopha* Nicaragua, and *I. i. iguana* Panama. *P*-values are HY-FDR adjusted. Non-significant results are indicated by n.s. These results exclude *I. delicatissima* and the only reported *I. delicatissima* x *I. iguana* hybrid (van Wagensveld & van den Burg 2018) as also in Figure 2A. SAB88 was also not included.

|  | Saba (native) | Saba (hybrid) | *I. i. rhinolopha*<br>Nicaragua |
| --- | --- | --- | --- |
| Saba (native) |  |  |  |
| Saba (hybrid) | 0.0017 |  |  |
| *I. i. rhinolopha*<br>Nicaragua | 0.073 (n.s.) | 0.0065 |  |
| *I. i. iguana* Panama | 0.0017 | 0.073 (n.s.) | 0.016 |

Considering comparisons of total size, ANOVA results indicated significant differences of mean SVL-corrected tail lengths among native Saba iguanas (*n* = 25), hybrid iguanas from Saba (*n* = 17), and non-native iguanas from St. Maarten (*n* = 22); *p* = 0.0312 (Figure 3). Tukey posthoc test showed only a significant difference between native Saba and non-native St. Maarten iguanas (*p* = 0.0265), and non-significant differences between Saba hybrids and the other two groups (*p* > 0.29).

**Figure 3.**
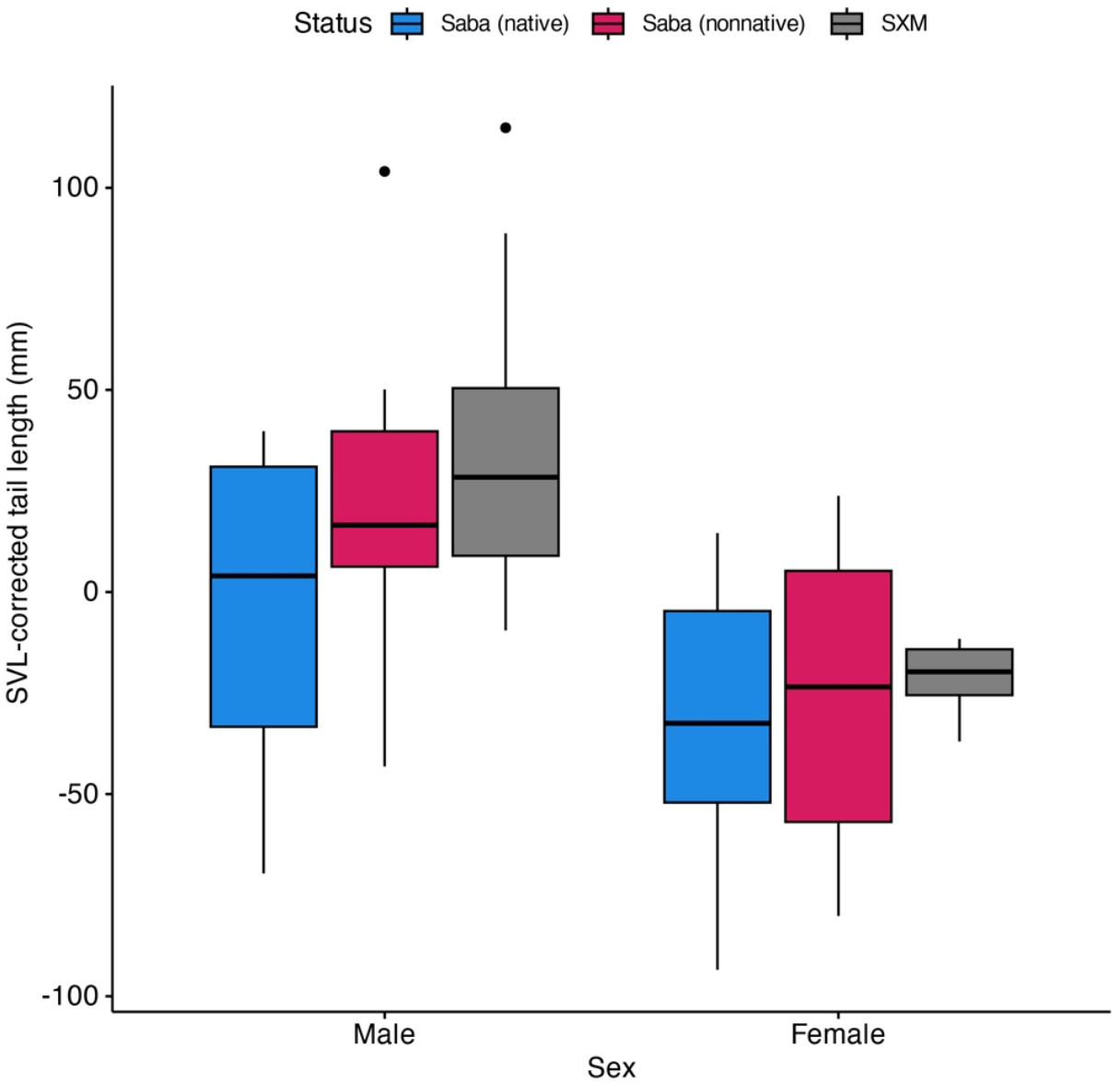
Boxplots showing differences between SVL-corrected tail length (overall size) between native Saba iguanas, hybrid iguanas from Saba, and non-native iguanas on St. Maarten.

## Discussion

In light of the ongoing threat of hybridization and genetic swamping by NNGI to native *Iguana* populations in the Lesser Antilles, clutch size has remained an unstudied life-history trait. Here we provide a first preliminary report on clutch size difference between native Saba iguanas, hybrid iguanas from Saba, and two mainland *Iguana* populations (from different mtDNA clades, II and III; see van den Burg et al. 2026). First, our results indicate that some mainland populations lay larger clutches compared to native Saba iguanas. This result aligns with the findings that insular lizards generally have smaller clutch size than to continental lizard populations (Novosolov and Meiri 2013; Meiri et al. 2020). Importantly, although using a small dataset, we find that hybrid iguanas on Saba appear to produce larger clutches than native Saba iguanas with a similar snout-vent length, which may facilitate their genetic swamping of native *Iguana* populations as witnessed throughout the Lesser Antilles.

Our NNGI removal campaign targets all sighted hybrid and nonnative iguanas on Saba, which appear to be unequivocally identifiable ater about two years of age (van den Burg et al. 2023, 2025b, unpublished data), when their detectability stabilizes to that of other adult animals. The clutch size data were, necessarily of an opportunistic nature, since the aim of the campaign is complete removal of all hybrid and non-native iguanas, preferably before they successfully nest. Data opportunities were additionally constrained by the limited number of hybrid iguanas due the invasion process having started less than 10 years ago (van den Burg et al. 2023, 2025b, 2025c, 2025d, unpublished data). This resulted in a relative small clutch size dataset, for which data could furthermore only be collected from adult females that successfully copulated and were removed during January-early March.

Clutch size in *Iguana* shows a high variation among sampled populations (e.g., Blankenship 1990; Nosolov et al. 2012; Knapp et al. 2016; Bock et al. 2022 and references therein). Presumably clutch size is at least partially affected by local climate and environmental conditions similar to other reptiles (e.g., Diaz et al. 2012; Roitberg et al. 2013), as well as island-mainland differences (Silliceo & Díaz 2010). There are similar signs from within *Iguana* (e.g., van Marken Lichtenbelt & Albers 1993; Knapp et al. 2016), but no study has of yet placed a narrow focus on this topic. Considering our case study, non-native iguanas on Saba originate from St. Maarten/Martin (van den Burg et al. 2018b, 2023), where a wide-spread non-native iguana population is present that has a genetic origin comprised of at least three of the four major mitochondrial clades within the *Iguana iguana* species complex (Clade I, II, and III; see van den Burg et al. 2026). Equally, our generated genetic data from hybrid and non-native individuals on Saba indicates similar genetic origins (Table 1 and Appendix 1). Currently there are no data on clutch sizes from non-native iguanas on St. Maarten/Martin, which would be interesting to better interpret the clutch size differences in our dataset. Considering clutch sizes of the three major mtDNA clades, although there likely is ample variation within each clade, our dataset includes data from Clades II and III (Fitch & Henderson 1977; Rand 1984; Miller 1987), but not from Clade I (van Marken Lichtenbelt & Albers 1993 did not provide the raw data). However, van Marken Lichtenbelt & Albers (1993) compared their SVL-clutch size data to those of Rand (1984), finding that clutch size for equal SVLs was significantly lower for Clade I compared to Clade II (see table 2 in van Marken Lichtenbelt & Albers 1993). Their assessment combined with our comparison of SVL-clutch size residuals (Table 4) suggests hybrid iguanas on Saba lay more eggs compared to native Saba iguanas as well as iguanas from Nicaragua and Curaçao, but not Panama despite a potential trend (*p* = 0.073). Clutch size data combined with genomic approaches could shed light on genes involved in fecundity and selective pressure in (hybrid) iguanas, as well as potential novel gene complexes and local adaptation in (the mostly mixed-origin) populations of NNGIs across the Lesser Antilles (Martin et al. 2015; Vuillaume et al. 2015; van den Burg et al. 2018b, 2023, 2024; Pounder et al. 2020).

For hybridization to occur, mating periods of native and non-native iguana should at least partially overlap. Equally variable compared to clutch size is the timing of reproduction within both *I. delicatissima* and *I. iguana* that follows local annual climate patterns (van den Burg et al. 2018a; Breuil et al. 2019). On Saba, we found that reproductive timing was similar between non-native/hybrid and native Saba iguanas, which is in contrast to some Lesser Antillean islands native to

*I. delica,ssima* (Breuil et al. 2019). However, since non-native iguanas from different geographic origins are present within the region and also hybridize (see Table 1; e.g., Vuillaume et al. 2015; van den Burg et al. 2023), future assessments of reproductive timing and geographic origin of non-native populations should be considered (e.g., on St. Martin), as well as potential changes in reproductive timing when NNGI arrive (plasticity) and when hybridization results in hybrid swarms. Understanding these population characteristics could inform tailored conservation strategies to bolster native populations against the threats posed by non-native iguanas, especially when targeting gravid females (with potential hybrid clutches) at nesting sites.

In conclusion, our results suggest that the observed decline in native *Iguana* populations is at least partially driven by genetic swamping, not only by a larger clutch size of pure NNGIs, but also through larger clutch size of hybrid iguanas compared to native iguanas (i.e., hybrid vigor). Because native iguanas can live to 24 years in the wild (Warret Rodrigues et al. 2021), and thus can remain present long enough ater incursions and establishment of NNGIs for (genetic) identification to make it possible to safeguard their genetic lineage for conservation purposes. We emphasize that despite the low variation in mtDNA across the range of *I. delica,ssima* (Martin et al. 2015), nuclear markers do indicate genetic structure among the different island populations (van den Burg et al. 2021). Since the maintenance of intraspecific genetic diversity is critical to long-term conservation of a species (e.g., Lacy 1997), identifying and capturing remaining native iguanas in hybridized and introgressed populations is critical for native *Iguana* conservation. We therefore urge regional partners to identify remaining native *Iguana* survivors within areas that have been heavily impacted by hybridization and introgression with non-native green iguanas (e.g., Grenada, Guadeloupe, and Martinique; Angin 2017; Breuil et al. 2019) so that their protection can be improved.

## Acknowledgments

We thank Peter Kuperus for performing all laboratory procedures and analyses, and the Public Entity Saba and Saba Conservation Foundation for on-island logistic support. We thank Charles Knapp for providing feedback and recommendations that greatly improved an earlier version of this manuscript. We also remain indebted to four anonymous reviewers for their critical feedback that allowed essential improvements to our manuscript. We are grateful for the financial support that allowed us to perform this study from the Dutch Iguana Foundation, the Iguanaland Conservation Fund, the Public Entity Saba within their invasive species project, and the Ministry of Agriculture, Fisheries, Food Security and Nature (LVVN) through the Wageningen University BO research program (BO-43-117-006) under Wageningen University and Research project number 4318100346-1. Fieldwork on Saba was approved under permits BC663.21, BC515.2023 and BC428.2024.

## Appendix

**Appendix 1**. Collected data on snout-vent length (SVL) and tail length from iguanas on Saba, including genetic assignment for mitochondrial (mtDNA; one haplotype), nuclear (nDNA; two haplotypes), and microsatellite (msat) data. Unless native, clade numbers in the assignment columns refer to mtDNA clades within the *Iguana iguana* species complex following van den Burg et al. (2026). Asterisk indicate data published in van den Burg et al. (2023). Native ND4 and MLH3 haplotypes correspond to HM352505 and Oti556119, respectively. All individuals had complete tails.

| Individual | Fate | SVL (mm) | Tail (mm) | mtDNA | nDNA |
| --- | --- | --- | --- | --- | --- |
| SAB01 (M)* | Alive | 230 | 595 | native | native/native |
| SAB03 (M)* | Alive | 405 | 1022 | native | native/native |
| SAB11 (F)* | Alive | 340 | 821 | native | native/native |
| SAB15 (M)* | Alive | 389 | 959 | native | native/native |
| SAB17 (F)* | Alive | 350 | 828 | native | native/native |
| SAB18 (F)* | Alive | 326 | 795 | native | native/native |
| SAB22 (M)* | Alive | 310 | 725 | native | native/native |
| SAB23 (F)* | Alive | 267 | 697 | native | native/native |
| SAB24 (M)* | Alive | 265 | 708 | native | native/native |
| SAB28 (M)* | Alive | 401 | 965 | native | native/native |
| SAB30 (F)* | Alive | 316 | 776 | native | native/native |
| SAB31 (F)* | Alive | 300 | 774 | native | native/native |
| SAB41 (F)* | Alive | 362 | 846 | native | native/native |
| SAB44 (M)* | Alive | 305 | 757 | native | native/native |
| SAB51 (F)* | Alive | 363 | 810 | native | native/native |
| SAB60 (F) | Alive | 361 | 838 | native | native/native |
| SAB61 (M) | Alive | 391 | 988 | native | native/native |
| SAB70 (F) | Alive | 357 | 902 | native | native/native |
| SAB75 (M) | Alive | 311 | 830 | native | native/native |
| SAB76 (M) | Alive | 371 | 957 | native | native/native |
| SAB82 (M) | Dead | 340 | 896 | native | native/native |
| SAB83 (M) | Alive | 383 | 907 | native | native/native |
| SAB84 (M) | Alive | 327 | 844 | native | native/native |
| SAB112 (M) | Dead | 271 | 675 | native | native/native |
| SAB114 (F) | Dead | 381 | 955 | native | native/native |
| SAB08 (M)* | Dead | 345 | 888 | Clade III: HM352508 | native/Clade III: OQ556118 |
| SAB57 (F)* | Dead | 359 | 838 | Clade III: HM352508 | native/Clade III: OQ556117 |
| SAB63 (M) | Dead | 410 | 1104 | Clade I: OQ556120 | native/ Clade I: KX610637_a |
| SAB69 (M) | Dead | 267 | 697 | Clade I: OQ556120 | native/ Clade I: KX610637_a |
| SAB80 (F) | Dead | 361 | 875 | native | native/Clade III: PV974928 |
| SAB85 (M) | Dead | 440 | 1073 | Clade I: HM352499 | native/Clade II: PV974929 |
| SAB87 (M) | Dead | 400 | 991 | Clade III: HM352508 | native/Clade III: OQ556117 |
| SAB90 (M) | Dead | 312 | 835 | Clade III: HM352508 | native/Clade III: OQ556118 |
| SAB91 (F) | Dead | 311 | 802 | Clade III: HM352508 | native/native |
| SAB94 (M) | Dead | 315 | 855 | native | native/Clade III: OQ556117 |
| SAB97 (F) | Dead | 357 | 811 | Clade III: HM352508 | native/Clade III: OQ556118 |
| SAB100 (F) | Dead | 369 | 921 | native | native/Clade I: OQ556119 |
| SAB102 (F) | Dead | 340 | 880 | native | native/Clade III: PV974928 |
| SAB105 (F) | Dead | 255 | 659 | Clade III: HM352508 | native/native |
| SAB115 (M) | Dead | 189 | 503 | Clade III: HM352508 | native/native |
| SAB116 (F) | Dead | 259 | 633 | Clade III: HM352508 | native/Clade III: OQ556117 |
| SAB117 (F) | Dead | 345 | 843 | native | native/Clade III: OQ556117 |

