## Supplementary figures for "Larger hybrid clutch size could drive regional displacement of native *Iguana* populations across the Lesser Antilles"

**Supplementary figure 1.** Results of quality of model fit evaluation for the generalized linear model excluding the bioclimatic variables.

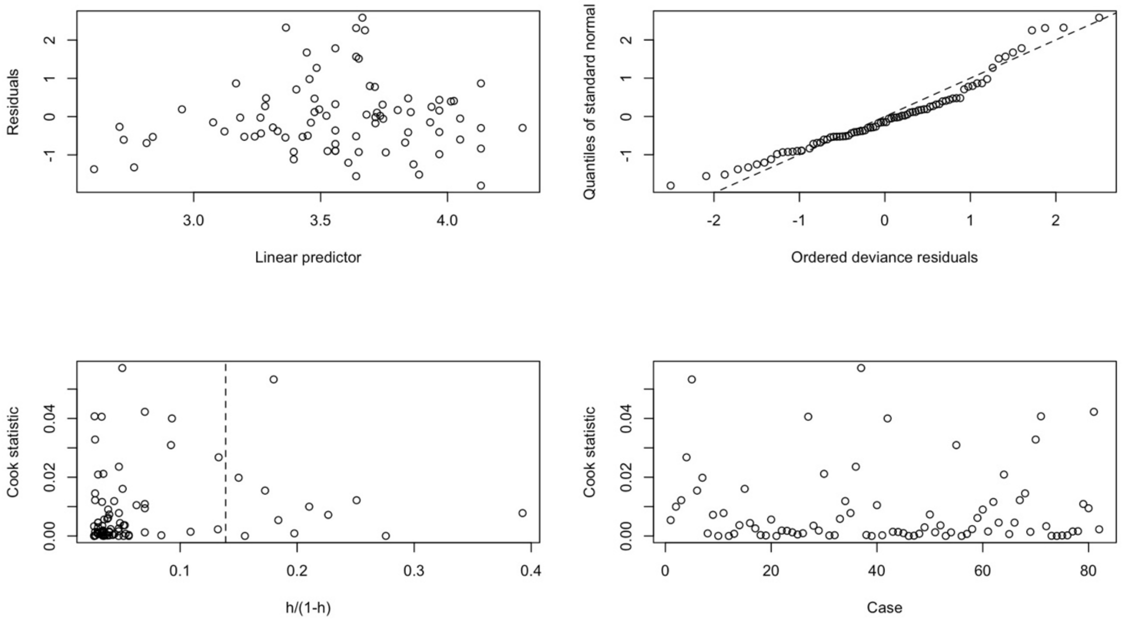


**Supplementary figure 2.** Results of quality of model fit evaluation for the generalized linear model including the bioclimatic variables.

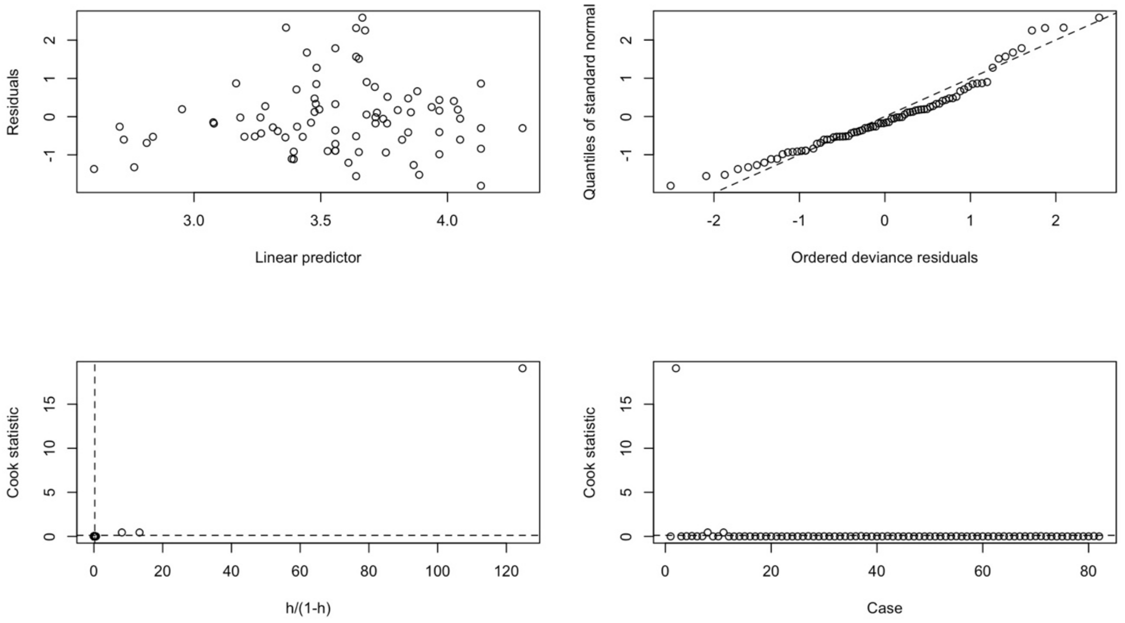
